# Medication-related osteonecrosis of the jaws after tooth extraction in senescent female mice treated with zoledronic acid: microtomographic, histological and immunohistochemical characterization

**DOI:** 10.1101/574111

**Authors:** Claudia Cristina Biguetti, André Hergesel De Oliva, Kent Healy, Ramez Hassan Mahmoud, Isabela Do Carmo Custódio, Dulce Helena Constantino, Edilson Ervolino, Marco Antonio Hungaro Duarte, Walid D. Fakhouri, Mariza Akemi Matsumoto

**Affiliations:** Department of Basic Sciences, São Paulo State University (UNESP) - School of Dentistry, Araçatuba - SP, Brazil; Department of Diagnostic and Biomedical Sciences, University of Texas Health Science Center at Houston, Houston-TX, United States; Department of Health Sciences Universidade Sagrado Coração – USC, Bauru – SP, Brazil; Department of Endodontics, São Paulo University – FOB/USP, Bauru - SP, Brazil

## Abstract

Treatment with cumulative dosages of zoledronic acid (ZA) in elderly patients is a risk factor for the development of medication-related osteonecrosis of the jaws (MRONJ), mainly related to surgical triggers such as tooth extraction. However, animal models for the investigation and understanding of MRONJ pathophysiology in senescent and postmenopausal stages remains to be developed and characterized. The aim of this study was to analyze MRONJ development in senescent female mice treated with cumulative dosages of ZA. For this purpose, twenty 129/Sv female mice, 64 weeks old, were treated with 0.9% saline solution as Control group (n=10), and with ZA at 250µg/Kg (n=10), once a week, starting 4 weeks before the upper right incisor extraction and until the end of the experimental time points (7 days and 21 days). At 7 and 21 days, specimens were harvested for microCT, histological, birefringence and immunohistochemical analysis. Clinically, an incomplete epithelialization was observed in ZA group at 7 days and a delayed bone matrix mineralization and collagen maturation at 7 and 21 days compared to the controls. Controls revealed sockets filled with mature bone at 21 days as observed by microCT and birefringence, while ZA group presented delayed bone deposition at 7 and 21 days, as well increased leukocyte infiltration and blood clot at 7 days, and increased bone sequestrum and empty osteocyte lacunae at 21 days (p<0.05). Also, ZA group presented decreased quantity TGFb+ and Runx-2+ cells at 7 days, and decreased quantity of TRAP+ osteoclasts compared to the control at 21 days (p<0.05). Togheter, these data demonstrate the usefulness of this model to understanding the pathophysiology of MRONJ.

## Introduction

The chronic use of antiresorptive drugs, such as nitrogen-containing bisphosphonates (nBP’s), is a risk factor in medication-related osteonecrosis of the jaw (MRONJ), and has been clearly associated with the disease onset [1–3]. MRONJ is a pathological condition characterized by a non-viable exposed maxillary or mandibular bone in the oral cavity in patients with previous or ongoing antiresorptive therapy with the exception of previous radiation treatment [1]. Risk factors of MRONJ include invasive dental procedures involving bone injuries, whereby dental extraction is the most common triggering factor [4, 5]. Furthermore, other procedures that require bone manipulation, such as dental implants, have a comparable risk to that of dental extractions [2, 6]. Additional comorbidities and systemic conditions have been reported as frequently associated with risk of developing MRONJ (i.e., cancer, advanced age, and the use of immune suppressive medications) [1, 2]. Other predisposing risk factors include drug dosage [7], potency, route of administration, and duration of nBP treatment [1, 6]. Notably, zoledronic acid (ZA) is 100 to 1000-fold more potent than other nBP’s [8], and therefore it is the first choice for the treatment of patient with bony metastasis [9]. It is also a highly effective drug for severe postmenopausal osteoporosis in elderly women [10], thereby reducing the risk of vertebral and hip fractures by up to 70% and 41%, respectively [11].

While MRONJ has well-defined histopathological characteristics, its pathophysiological mechanism remains to be fully elucidated [1]. MRONJ human biopsies exhibit empty osteocyte lacunae, lack of osteoblasts along new bone, and detached osteoclasts. MRONJ biopsy samples are frequently associated with a secondary infection [12, 13], and may exhibit similar histological characteristics to osteomyelitis and osteoradionecrosis [13]. Previous studies have explored MRONJ animal models to provide explanations for the disease process. Hypotheses examined in these studies include over-suppression of osteoclasts bone resorption in the jaws, more than in endochondral bones [14], angiogenesis inhibition [15, 16] and more recently, alterations of innate and adaptive immune responses [17, 18].

It is important to emphasize that bone healing depends on the initial inflammatory response, which can be affected by the type drug and the compromised bone cells in the injured areas [19]. Therefore, an immunological mechanism involved in the development of MRONJ is an important process that needs further investigation using a suitable animal model. Mice are considered a reliable *in vivo* system for immunological investigations and have conferred multiple advantages over other animal models therein [20]. Namely, the small size of the animal requires reduced amount of the drug, rapid cumulative effects due to rapid mice metabolism, and the possibility of using genetically-manipulated animals to determine a cause-and-effect relationship between the target disruption of a gene and its function [21–23]. Several MRONJ models have been developed using mainly C57BL/6 and other mice strains (S1 Table), [15, 17, 23–39]. The C57BL/6 is the most widely used inbred strain and with a large source of genetically modified lines [40]. However, it has overall less bone volume than other strains, such as the 129Sv [41]. Therefore, 129Sv could provide an advantage on C57Bl/6, as a mouse strain with more robust bone for studying conditions where bone density is an important variable.

The majority of previous studies reported the use of ZA via IP (intraperitoneal) or IV (intravenous) administration as an antiresorptive medication of choice for inducing MRONJ lesions in mice [15, 17, 24, 25, 27, 29, 30, 32, 33, 35, 37, 39, 42–46], considering its higher potency when compared to other IV or oral nBP’s [47]. Moreover, all mouse studies presented in S1 Table were performed in young or adult animals (age varying from 6 to 16 weeks old). The age of the tested mice is an important variable that possibly increases the risk for MRONJ onset. According to the last clinical update from AAOMS [3] the risk of developing MRONJ among osteoporotic patients exposed to nBPs remains very low, but affects mainly the population over the age of 55. The average age of menopaused women is 51 years old, and they can live 30% of their life in a postmenopausal state, considering a life expectancy of 80 years old approximately [48]. Considering these epidemiological and clinical data, senescent female mice could provide a suitable model to investigate bone healing after dental procedures in this treatment conditions, using ZA or other MRONJ potential drugs. Thus, the aim of this study is to analyze bone healing after tooth extraction in senescent mice treated with cumulative doses of ZA, and investigate whether 129Sv senescent female mice are appropriate model by inducing and understanding the MRONJ development via microtomographic, histopathological and immunohistochemical characterization.

## Material and Methods

### In vivo assays

In vivo assays were performed by using a total of twenty 129 S/v female mice (60-weeks-old), provided by the animal facilities of University of Sao Paulo (University of São Paulo, Ribeirão Preto, São Paulo, Brazil) and maintained in the animal facilities of the Universidade Sagrado Coração, Bauru, São Paulo, Brazil). Animals were handled according to the recommendations in the Guide for the Care and Use of Laboratory Animals of the National Institutes of Health (Institute of Laboratory Animal Resources (U.S.). The experimental protocol was performed according to ARRIVE guidelines [49] and approved by the local Institutional Committee for Animal Care and Use of the Universidade Sagrado Coração (CEUA Protocol number #9589271017). First, estrus cycle was evaluated by vaginal cytology following previous recommendation and criteria for stained vaginal smears with Toluidine blue O stain [50, 51]. Sixty-weeks old mice presented cytology with predominance of leukocytes and few epithelial cells for four consecutive weeks, indicating a persistent diestrum and were considered in the postmenopausal period. At 64 weeks old, the mice were divided into control (n=10) and experimental (n=10). The Control group was treated with IP injection of 0.01ml of sterile 0.9% saline solution; while the ZA group was treated with an IP injection of Zoledronic Acid (Merck KGaA, SML0223, Darmstadt, Germany) at 250 µg/kg diluted in 0.9% sterile solution once a week. Mice received IP injections of ZA once a week, starting 4 weeks before tooth extraction and continuing until the end of experimental periods (7 days or 21 days). Animals were fed with sterile standard solid mice chow (Nuvital, Curitiba, PR, Brazil) and sterile water, except on the first 72 hours after surgery, in which diet was soft for postoperative recovery and to facilitate the ingestion of food. Animals were clinically monitored and controlled regarding body loss, feeding and behavior. No antibiotics or anti-inflammatory drugs were administered to the animals after tooth extraction, in order to avoid interference with investigated inflammatory pathways [22]. experimental groups were comprised of 5 animals per group/time point (7 and 21 days), and specimens were harvested for microCT, histological, birefringence and immunohistochemical analysis.

### Mice tooth extraction model

Tooth extractions were performed as previously described [52, 53]. Animals received general anaesthesia by intramuscular administration of ketamine chloride (80 mg/kg) (Dopalen, Agribrans do Brasil LTDA, SP, Brazil) and xylazine chloride (15 mg/kg) (Anasedan, Agribrands do Brasil LTDA, SP, Brazil). Each mouse’s oral cavity was divided into two parts: upper right side with extraction (E side), and upper left side without extraction (NE side). Then, the right upper incisor luxation was performed with a dental exploratory probe until the tooth became mobile, and was then removed delicately with forceps. At the end of experimental time points, mice were euthanized, and maxillae containing the socket area were harvested. Additionally, femur and L5 vertebrae were collected for supplementary analysis at the experimental period of 7 days (after 5 doses of ZA). Bone specimens were immediately fixed in PBS-buffered formalin (10%) solution (pH 7.4) for 48h at room temperature, subsequently washed overnight in running water and maintained temporarily in alcohol fixative (70% hydrous ethanol) and scanned for μCT analysis, and then decalcified in 4.13% EDTA (pH 7,2) for histological processing.

### Micro CT analyses

Bone specimens were scanned by the Skyscan 1174 System (Skyscan, Kontich, Belgium) at 50 kV, 800 μA, with a 0.5 mm aluminium filter, 180 degrees of rotation and exposure range of 1 degree and a resolution of 14μm pixel size. Subsequently, images were reconstructed using the NRecon software, following three-dimensional (3D) reconstruction of the images using CTVox, and the quantitative parameters were assessed using CTAn software. For the quantitative analyses, guidelines of μCT characterization were used as previously described [54]. The NE side maxilla and E side for the newly bone formed in the extracted alveolar area were analyzed following microtomographic parameters reported in previous studies [52]. Briefly, fraction of bone volume (BV/TV, %) was measured considering the region of the alveolar process of the upper incisor facing the animal palate, in a region of interest (ROI) of 1mm length and a diameter of 0.9 mm, as demonstrated later in the results. For the newly bone formed in the extracted alveolar area, BV/TV (%) was measured in a segmented in a cylindrical ROI covering the entire length of the alveolus (3mm) and a diameter of 1 mm.

### Histology sample preparation

Maxillary bone specimens were immersed in buffered 4% EDTA for demineralization for 3 weeks, then the specimens underwent histological processing for embedding in paraffin blocks. Transversal serial 5µm histological slices from coronal and medial third were cut for histological staining with H&E, modified Goldner’s Trichrome/Alcian Blue, and Picrosirius Red.

### Histological and histomorphometric analysis

A qualitative histopathological analysis was performed considering parameters of healing and MRONJ. Histomorphometric analysis was performed using a total of eight technical sections from middle regions of the alveolar socket stained by H&E. These samples were used to quantify the following components: blood clot, inflammatory infiltrate, blood vessels, osteocytes and empty lacunae. Quantification of histological parameters was performed by a single calibrated investigator with a binocular light microscope (Olympus Optical Co., Tokyo, Japan) using a 100× immersion objective. Eight histological fields per H&E stained section, comprising the region of alveolar socket were captured using a 100× immersion objective. A grid image was superimposed on each histological field, with ten parallel lines and 100 points in a quadrangular area, by using Image J software (Version 1.51, National Institutes of Health, Bethesda, Maryland, USA). Only the points coincident with the histological parameters were considered and the total number of points was obtained to calculate the area density for each healing component in each section [22].

### Modified Goldner’s Trichrome/Alcian Blue

Modified Goldner’s Trichrome/Alcian Blue (GT+ Alcian Blue) was used to stain newly formed bone matrix and cartilage with green and dark blue color, respectively. Hematoxylin was used as a counter stain for nuclei. Particularly, this stain offers a wider array of colors and contrast, for a clearly differentiation between the different inflammatory cells and bone components. Histological sections were incubated for 30 min at 56 °C, followed by washing in distilled water for 5 min. Then, the sections were stained with Alcian Blue for 15 minutes, and washed 3 times with distilled H_2_O for 3 min each time. Subsequently, sections were stained with Hematoxylin Harris for 45 seconds, rinsed in tap water for 6 minutes and distilled water for 5 minutes. Then, sections were stained in Fuchsin-Ponceau solution for 30 minutes, rinsed in 1.0% acetic acid solution for 1 minute, then stained in Orange G dye for 8 minutes, rinsed again in 1.0% acetic acid solution for 1 minute, stained in Light Green solution for 20 minutes, and finally rinsed in 1.0% acetic acid solution for 1 minute. Lastly, the slides were dehydrated in ascending ethanol solution followed by 2 washes in xylene for 3 min each time, and mounted with mounting media.

### Birefringence analysis for collagenous content maturation

For analyzing the quality and quantity of collagen, birefringence analysis was performed with Picrosirius-polarization method. For this method, eight histological sections from central region of the each alveolar socket were stained with Picrosirius Red stain and the images of 4 histological fields a 40× objective were captured by a polarizing lens coupled to a binocular inverted microscope (Leica DM IRB/E), as previously described [55]. Only socket area filled with new tissue was considered in this analysis. At the beginning, spectra of green, yellow and red colors were defined, followed by RGB (red, blue, green) values for each color spectrum and the quantity of pixels2 of each color correspondent to each field. Afterwards, the mean pixels2 values considering the intensity of birefringence brightness (pixels^2^) was performed using the AxioVision 4.8 software (Carl Zeiss) to define total area of green (thin and immature fibers), as well yellow and red collagen fibers (thicker and mature).

### Immunohistochemistry and Immunofluorescence

Bone sections from socket areas at 7 and 21 days post tooth extraction were deparaffinized following standard procedures. For immunohistochemistry, slices were pre-incubated with 3% hydrogen peroxidase block (Spring Bioscience Corporation, CA, USA) and subsequently incubated with 7% NFDM to block serum proteins. For the central regions of sockets, sections were incubated with goat polyclonal anti-Runx2 (sc8566, Santa Cruz Biotechnology, Carpinteria, USA), rabbit polyclonal anti-F4/80 (sc26643, Santa Cruz Biotechnology, Carpinteria, USA) at a concentration of 1:100, and rabbit polyclonal anti-TGF-β (sc7892, Santa Cruz Biotechnology, Carpinteria, USA), goat polyclonal anti-TRAP (sc30832, Santa Cruz Biotechnology, Carpinteria, USA), at a concentration of 1:200, for 1h at room temperature. For detection methods, universal immuno-enzyme polymer method was used and sections were incubated in immunohistochemical staining reagent for 30 min at room temperature. For detection of antigen–antibody we used 3-3’-diaminobenzidine (DAB), followed by counter-staining with Mayer’s hematoxylin. For control staining of the antibodies, serial sections were treated only with the Universal immuno-enzyme polymer, in a separate preparation. Immunofluorescence was performed following protocols previously described [56]. For detection of cell proliferation in the oral mucosa, sections from the initial portions of sockets were incubated with primary antibody for Proliferating Cell Nuclear Antigen (PCNA) (#MAB424, Merck Millipore, Darmstadt, Germany) diluted to 1:150 in blocking solution and incubated at 4°C overnight. After washing steps in 1×PBS (3 times, 10 min each wash), the sections where incubated with Cy3 secondary antibody (#715-165-150, Jackson ImmunoResearch Laboratories, West Grove, PA, USA), diluted to 1:200 in blocking solution, during 2 hours in a dark chamber at room temperature. Then the sections were washed 3 times in 1×PBS (10 min each wash) and the nuclei were counter-stained with DAPI (D9542-50, Sigma-Aldrich Corp., St. Louis, MO, USA) for 10 minutes of incubation in a dark chamber. After the last washing step in 1×PBS for 10 min, the slides were mounted with ProLong Gold Antifade Reagent (P36930, Life Technologies, Carlsbad, CA, USA).

### Quantification of TRAP, Runx-2 and F4/80 immunolabeled cells

The analysis of immunolabeled TRAP, Runx-2 and F4/80 was performed following the similar criteria described previously for histomorphometric analysis and as previously reported [53]. Briefly, a biological replicate of five samples for each experimental period and strains were used for quantitative analysis. A total of 3 sections of each sample containing the central region of the alveolar socket were used for targets quantification [53]. A total of eight fields were analyzed by using Image J software (Version 1.51, National Institutes of Health, Bethesda, Maryland, USA). Only the points coincident with the immunolabeled cells were considered in cell counting and the mean for each section was obtained for statistical analysis.

### Statistical analysis

Quantitative data were statistically analyzed for distribution of normality using Shapiro-Wilk and D’Agostino-Pearson omnibus test. For data with normal distribution, multiple comparisons among data were analyzed by One-Way analysis of variance (ANOVA) followed by the Tukey post test. For data that did not fit in the normal distribution, the Mann-Whitney and Kruskal-Wallis test were used followed by the Dunn’s test. The alpha level for all tests was set to (5%), then p<0.05 were considered statistically significant. Data sets are presented with descriptive statistics, containing mean and standard deviation. All statistical tests were performed using the GraphPad Prism 5.0 software (GraphPad Software Inc., San Diego, CA, USA).

## Results

Given the advantages for using a mouse as an animal model for immunological studies, our primary interest in this study was to investigate the bone healing in senescent and postmenopausal 129 Sv female mice treated with ZA, in order to provide a tool for understanding the MRONJ pathophysiology. Before we induced MRONJ through tooth extraction, we characterize the senescence and postmenopausal state in the female mice. The animals were evaluated for estrous cycle and presented a predominance of leukocytes and few epithelial cells for 4 consecutive weeks, indicating a persistent diestrum and a postmenopausal state (Fig. 1A). At 64 weeks old, Control and ZA group mice began receiving IP injections of each substance, 0.9% saline solution or ZA, respectively (Fig. 1B).

**Fig. 1.**
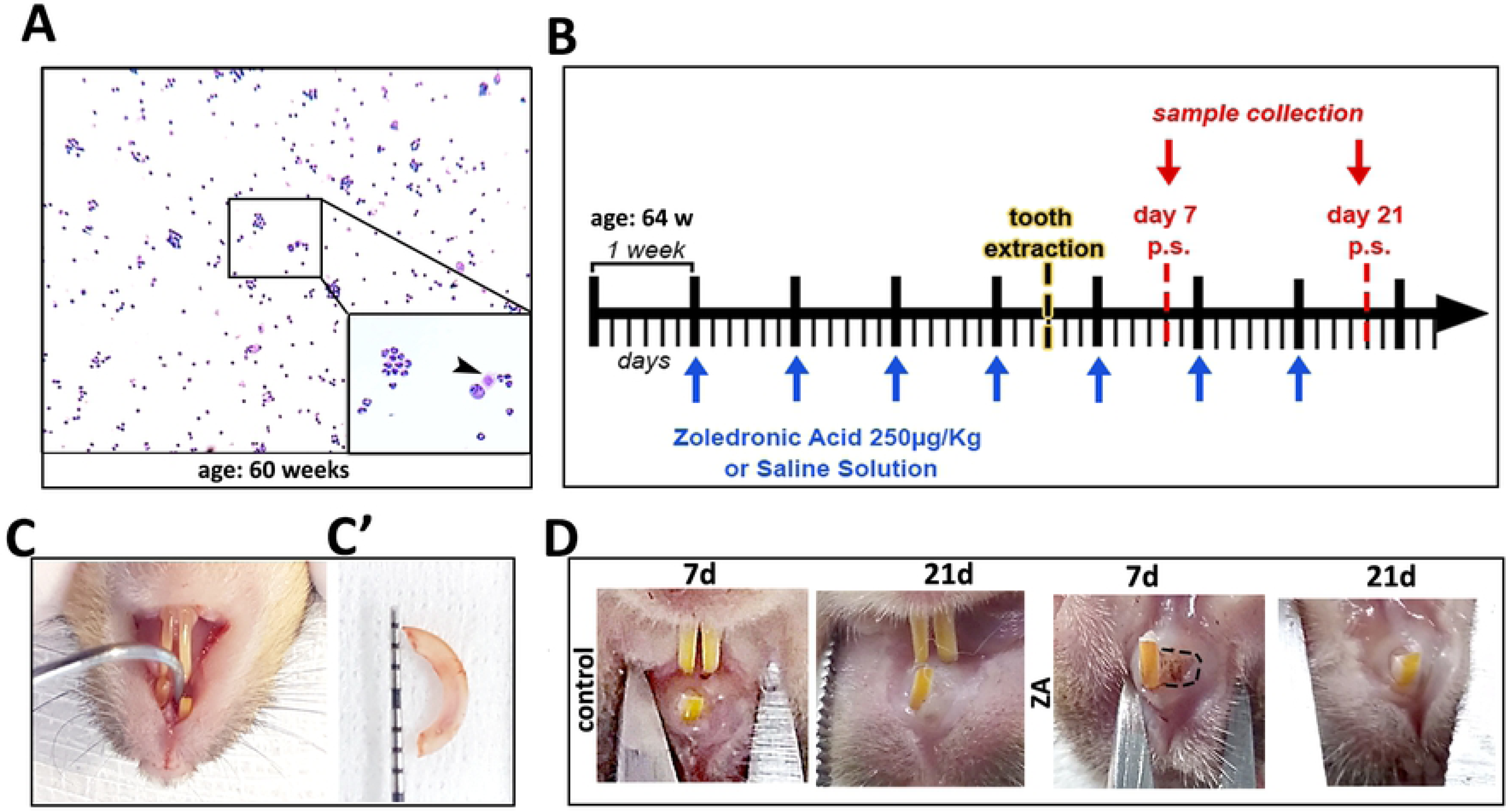
Experimental design of MRONJ model in senescent female 129 Sv mice. A) Vaginal smears with Toluidine blue O stain showing predominance of leucocytes and few nucleated epithelial cells (arrow head) for 4 consecutive weeks, as indicated for 60 weeks of age. B) Schematic timeline for Control and ZA groups: mice were treated with 0.9% saline solution or ZA at a dosage of 250µg/Kg, starting 4 weeks before tooth extraction. At 68 weeks of age, mice were submitted to atraumatically extraction (C) of right upper incisor (C’) and were euthanized at 7 and 21 days after the surgical procedure. D) Macroscopical occlusal views present clinical aspect of healing oral mucosal post tooth extraction:100% of control mice had a complete mucosal closure at 7d and 21 days, while 40% of ZA treated mice presented delayed epithelial socket closure at 7 days (dotted line), but also a complete mucosal closure at 21 days.

### Oral Mucosal Evaluation

At 68 weeks old, right upper incisor was atraumatically extracted four weeks after the initial IP injection with both solutions (Fig. 1C and C’). The animals were monitored daily for clinical post-operative symptoms or signs of infection or distress. At 7 days post-operation, 100% of Control group presented a complete healing of the oral mucosal, covering the surgical wound. On the other hand, 40% of ZA group clinically presented delayed epithelial socket closure. However, 100% of mice of Control and ZA group mice presented a complete mucosal closure at 21 days (Fig. 1D).

### Microtomographic Analysis of E site and NE site Maxillary Bone

After euthanasia, maxillae from 7 and 21 days post extraction were collected for initial microtomographic qualitative and quantitative analyses. Extant bone and new bone formation were analyzed at NE sites (Fig. 2A-A’) and E sites (Fig. 2B-B’), respectively. On the NE sites, the cumulative effects of ZA administration are evident in the significant increase in BV/TV of alveolar ridges in the ZA group compared to the Control group. On the NE sites, at 7 days post tooth extraction (or 5 injections of ZA or vehicle), the ZA group presented 86.67 ± 6.10% of BV/TV compared to 78.25 ± 9.74% of the Control group. At 21 days post tooth extraction (or 7 injections of ZA or vehicle), the ZA group presented 91.67 ± 1.52 % of BV/TV compared to 72.00 ± 16.10% of the Control group (Fig. 2A-A’). Regarding the newly formed bone in the alveolar sockets post tooth extraction (E sites), no significant differences were observed at 7-day time point between both groups. In comparison, both groups (Control and ZA) presented gradual bone apposition, with significant increased values of BV/TV compared to the initial time point at 21 days. In the comparison between groups, Control group presented significant increased bone formation compared to ZA, with 55.82 ± 3.51% vs. 44.45 ± 4.96 % (Fig. 2 B-C).

**Fig. 2.**
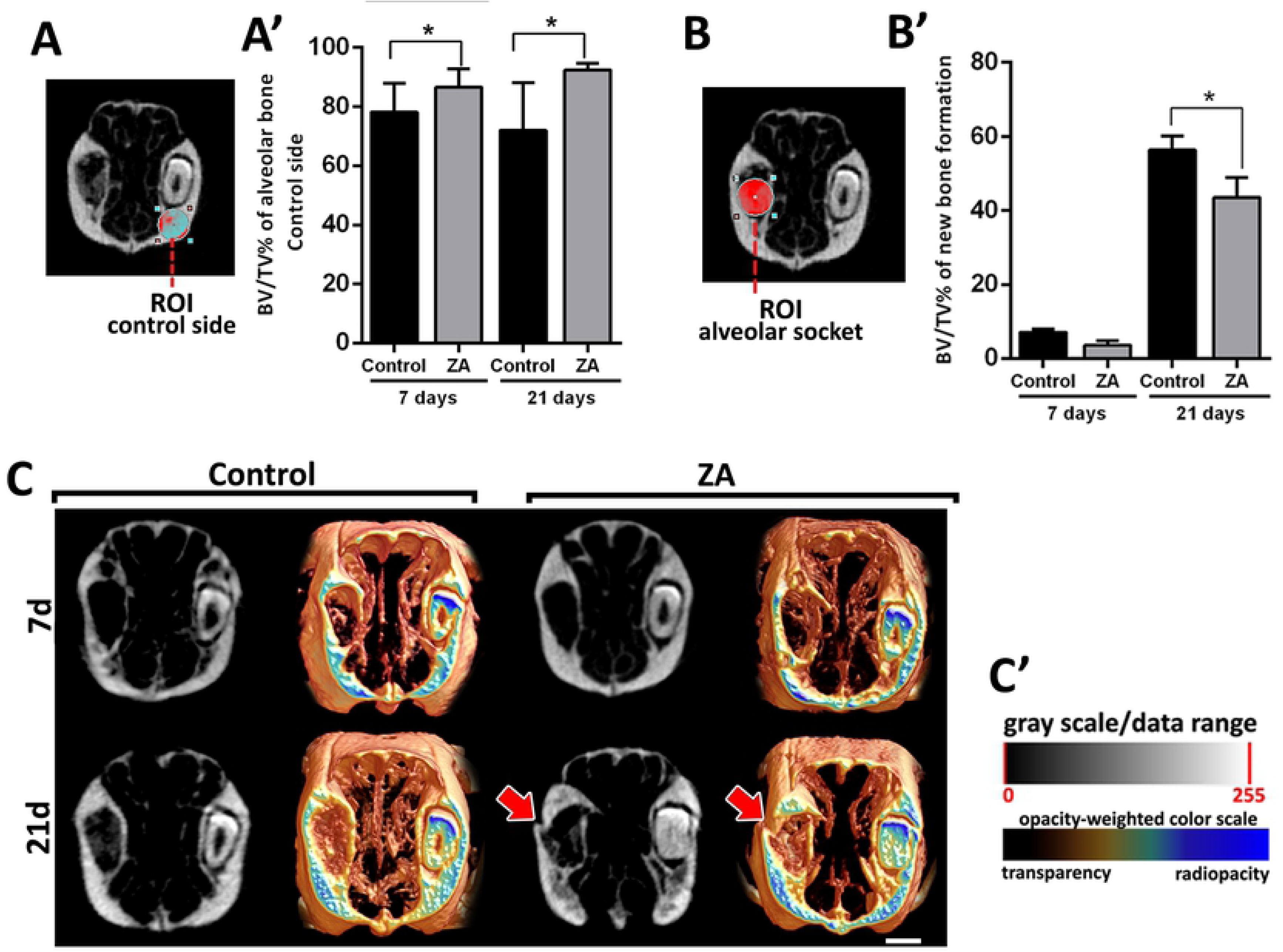
MicroCT evaluation of ZA cumulative effects on maxillary bone in NE sides and E sides (alveolar sockets post tooth extraction) in Control and ZA treated mice. Senescent 129 Sv-WT female received IP injections of 0.9% saline solution (Vehicle) or 250 µg/Kg once a week, and upper right incisor were removed at 4 week of each treatment. Mice were euthanized for maxillary bones removal after 7days and 21days post tooth extraction.A) BV/TV (%) of intact bone (NE side) and B) in the alveolar socket post tooth extraction (E side). C)Three-dimensional representative images obtained with the CT-Vox software (Skyscan, Kontich, Belgium), showing transverse sections of Control and ZA maxillary bones at 7days and 21days post tooth extraction. A bone fracture is observed in ZA treated mouse at 21 days post tooth extraction (red arrow). Results are presented as the means (±SD) of each parameter. Symbol *indicate a statistically significant difference vs control (p<0.05).

### Birefringence birefringence analysis for Collagen Evaluation

Using birefringence analysis, we evaluated the quality and quantity of collagen in new bone formed after tooth extraction in E sites (Fig. 3A-E). The transversal sections containing sites of healing (Control group) or MRONJ (ZA group) were evaluated for collagen fibers maturation, by the presence of birefringent collagen fibers (green, yellow and red) from the new bone or initial granulation tissue. While the Control group presented a gradual maturation of collagen fibers along 7 and 21 days, as well as a predominance of red spectrum fibers color at 21 days, the ZA group comparatively presented a lower quantity of collagen content at 7 days, and a predominance of green birefringent sprectrum collagen fibers at 21 days, indicating a disorganized organic matrix.

**Fig. 3.**
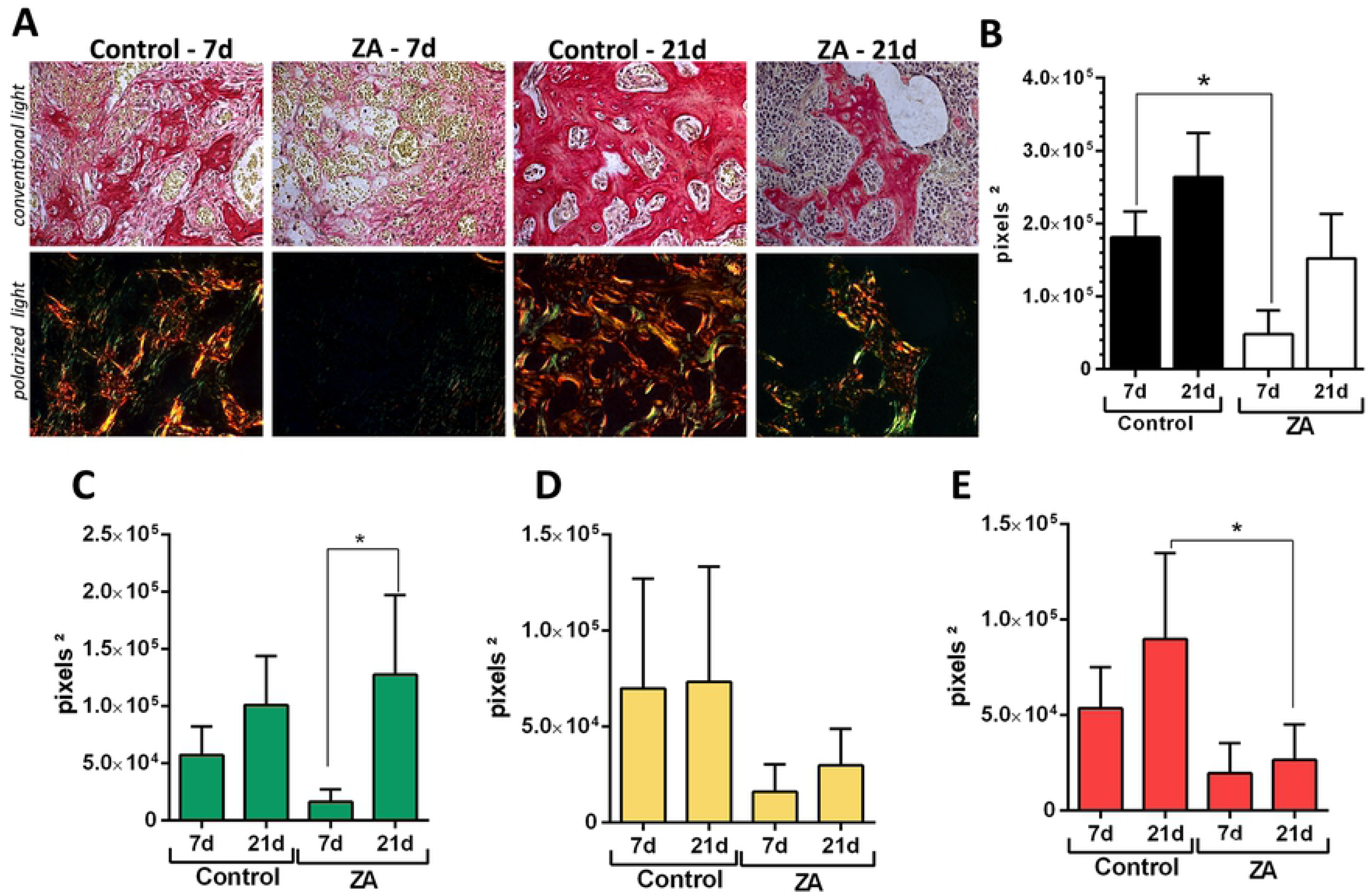
Birefringence analysis of collagen fibers in the alveolar sockets post tooth extraction (E sides) in Control and ZA treated mice. Senescent 129 Sv-WT femalemice received IP injections of 0.9% saline solution(Vehicle) or 250 µg/Kg once a week and upper right incisor were removed after 4 weeks of Vehicle or ZA treatments. Mice were euthanized for maxillary bones removal after 7days and 21days post tooth extraction.A)Representative transversal sections alveolar socket upon polarized and conventional light, to evaluate collagen fibers maturation in the different experimental groups and periods. As visualized upon polarized light, green birefringence color indicates thin fibers; yellow and red colors at birefringence analysis indicate thick collagen fibers. Original magnification 40x. Intensity of birefringence measured from Image-analysis software (AxioVision, v. 4.8, CarlZeiss) to identify and quantify total area of collagen fibers (B) and area of collagen from each birefringence color (pixels^2^) (C-green, D-yellow and E-red). Results are presented as mean and SD of pixels^2^ for each color in the birefringence analysis. Symbol * indicates a statistically significant difference *vs* control (p<0.05).

### Histopathological and Histomorphometric analysis

At day 7, dental sockets from the Control group healed without any complications, filled by a highly cellular and vascularized granulation tissue, when discrete osteogenesis was observed (Fig. 4). After 21 days, mature bone trabecullae could be seen within a well-organized medulla. In the ZA group at day 7, a delay of healing process was indicated by the presence of focus of blood clot and its persistence until 21 days, as also confirmed by histomorphometry (Fig. 4A-B). At day 21, intense leukocyte infiltration was observed, amongst thin primary trabeculae (Fig. 4A and C). Also, areas of non-viable bone were noted, but interestingly serving as a scaffold for new bone formation in some areas (Fig. 4A, D-E), along some discrete bone sequestrum. No biofilm formation was detected at this site of analysis for both the Control and ZA groups. In the ZA group, other findings were highlighted in the Fig. 5, such as nodular formation of mineralized matrix surrounding blood clot, several detached osteoclasts in the connective tissue, short newly formed bone trabeculae and non-viable alveolar bone presenting empty lacunae of osteocytes close to the remaining periodontal ligament. Interestingly, the NE side (containing the left upper incisor), presented alveolar bone containing filled osteocytes lacunae in both Control and ZA groups (S1 Fig).

**Fig. 4.**
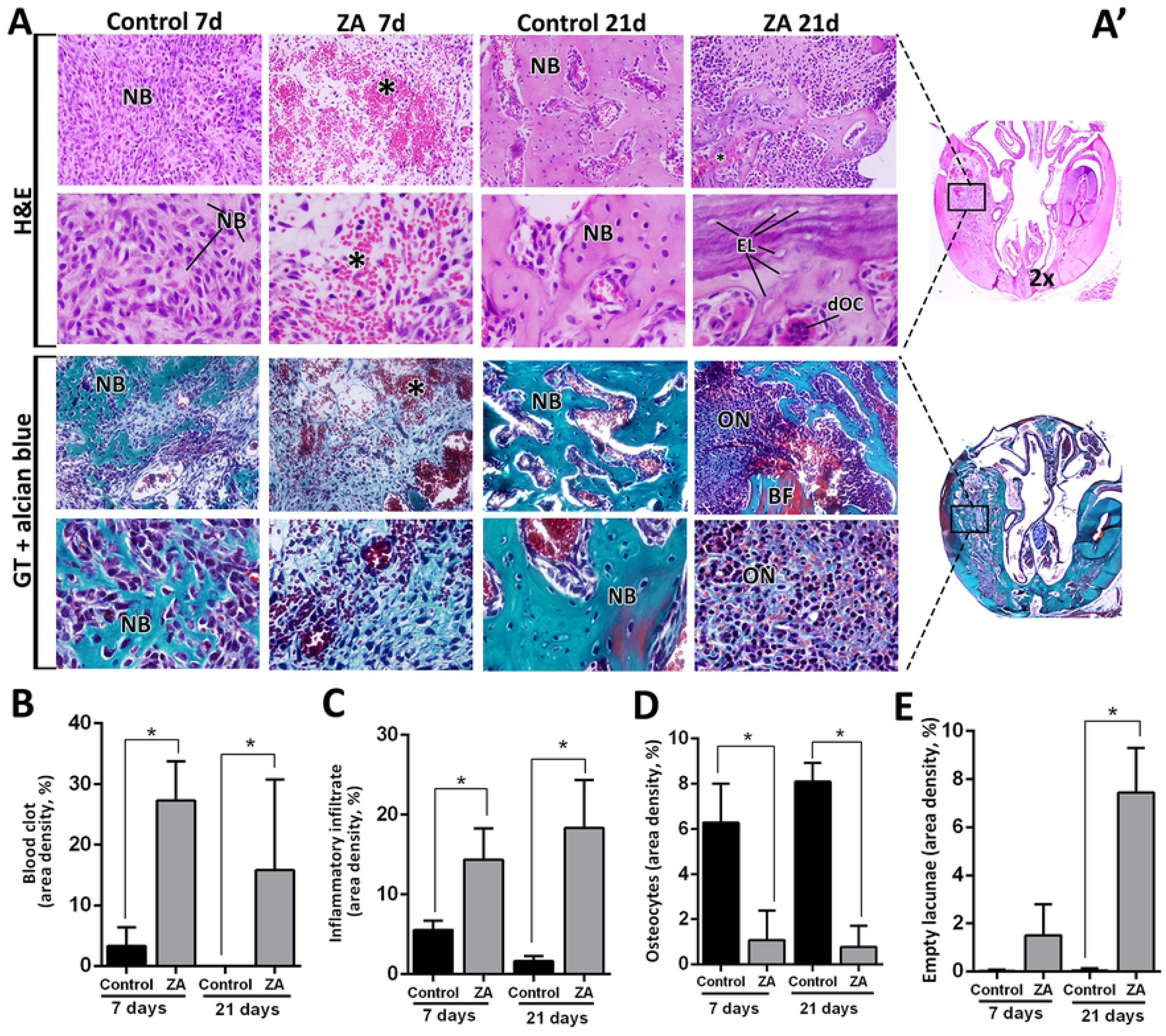
Histopathological and histomorphometric analysis of alveolar sockets post tooth extraction (E sides) in Control and ZA treated mice. Senescent 129 Sv-WT femalemice received IP injections of 0.9%saline solution(Vehicle) or 250 µg/Kg one a week and upper right incisor were removed after 4 weeks of Vehicle or ZA treatments. Mice were euthanized for maxillary bones removal after 7days and 21days post tooth extraction.A)Representative transversal sections are observed throughout days 7 and 21, from the central area of alveolar sockets (A’). Histological slides were stained with H&E (upper panel) and GT+Alcian blue (lower panel) and images were captured at 2x (entire section at left side) and 100x magnification (panels). Components of healing and MRONJ are indicated with letters and symbols, as follow: NB = new bone formation, ON = MRONJ sites with elevated inflammatory infiltrate, * = remaining/ non resorbed blood clot, BF = Bone fracture, EL= Empty Lacunae, dOC = detached OC. B-E) Histomorphometric analysis of healing or MRONJ parameters, including B) Remaining focus of blood clot, C) Inflammatory infiltrate, D) Osteocytes and E)Empty lacunae indicating non-viable bone. Results are presented as the means (±SD) of area density for each component. Symbol *indicate a statistically significant difference vs control (p<0.05).

**Fig. 5.**
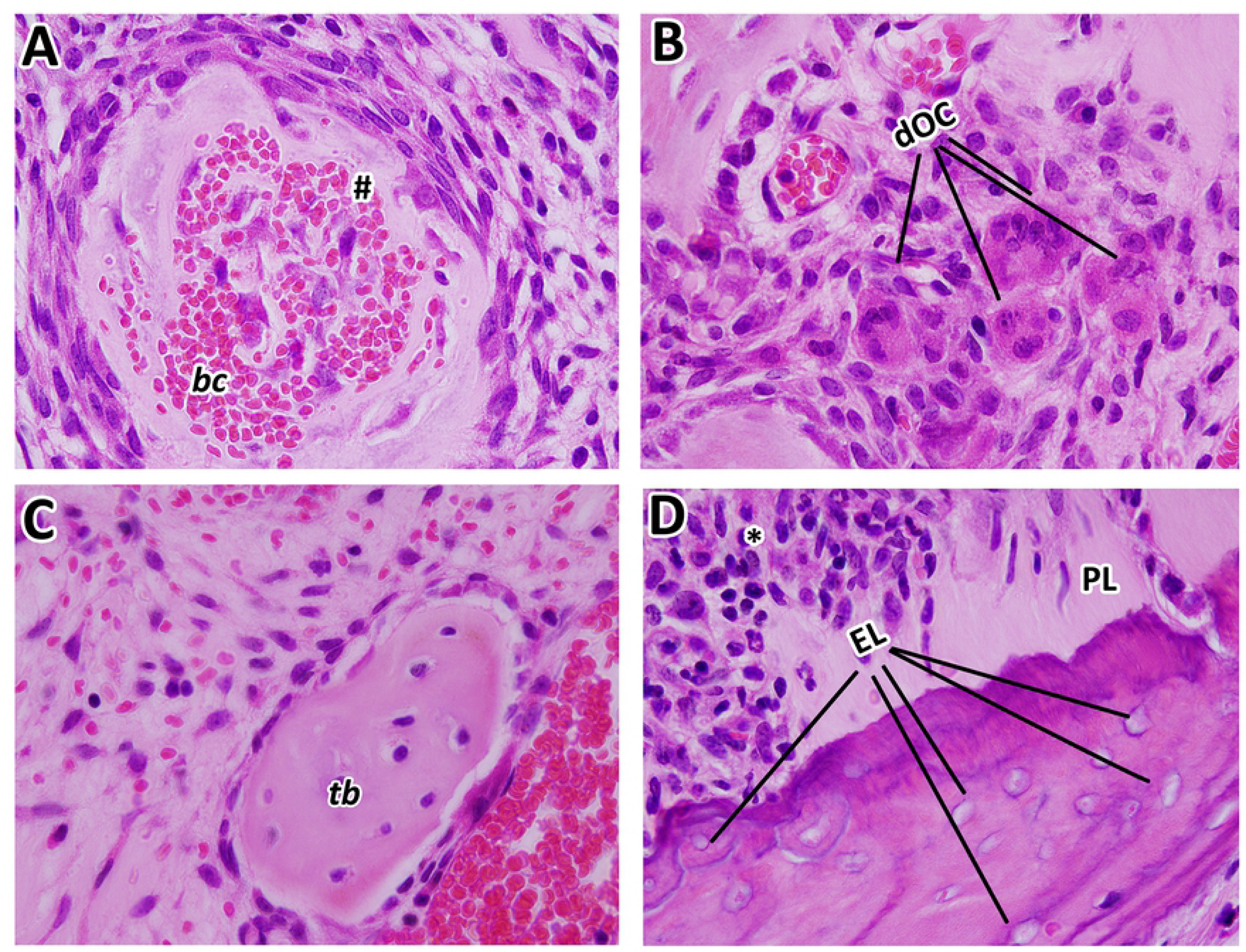
Details of MRONJ lesions in 129Sv senescent female mice treated with ZA at 21 days post tooth extraction. Representative transversal sections are observed throughout 21days, from the central area of alveolar sockets, 100x magnification. A) Nodular formation of mineralized matrix surrounding blood clot, associated with a highly cell contingent; B) detached osteoclast in the connective tissue; C) short newly formed bone trabeculae; D) non-vital alveolar bone presenting empty osteocytes lacunae close to the remaining periodontal ligament. MRONJ are indicated with letters and symbols, as follow: # nodular formation, dOC = detached OC, PL = remaining Periodontal Ligament, EL= Empty Lacunae, * = inflammatory infiltrate.

### TRAP+ Osteoclast Staining Analysis

Despite clear identification of some multinucleated OCs in the H&E stained sections, TRAP immunostaining was performed in alveolar sockets to accurately evaluate the area density (%) of OCs in the Control and ZA groups. TRAP+ OCs were found detached from bone, mainly in the ZA group (1.60 ± 0.81) compared to the Control (0.016 ± 0.28) at 21 days, while the Control group had an increased quantity of attached TRAP+ OCs (6.40 ± 2.45) compared to the ZA group (1.93 ± 0.50) in this same time point. In total, TRAP+ OCs were significantly reduced in ZA group (3.53 ± 0.64) compared to Control (6.56 ± 2.17) at 21 days (Fig. 6, Table 1).

**Fig. 6.**
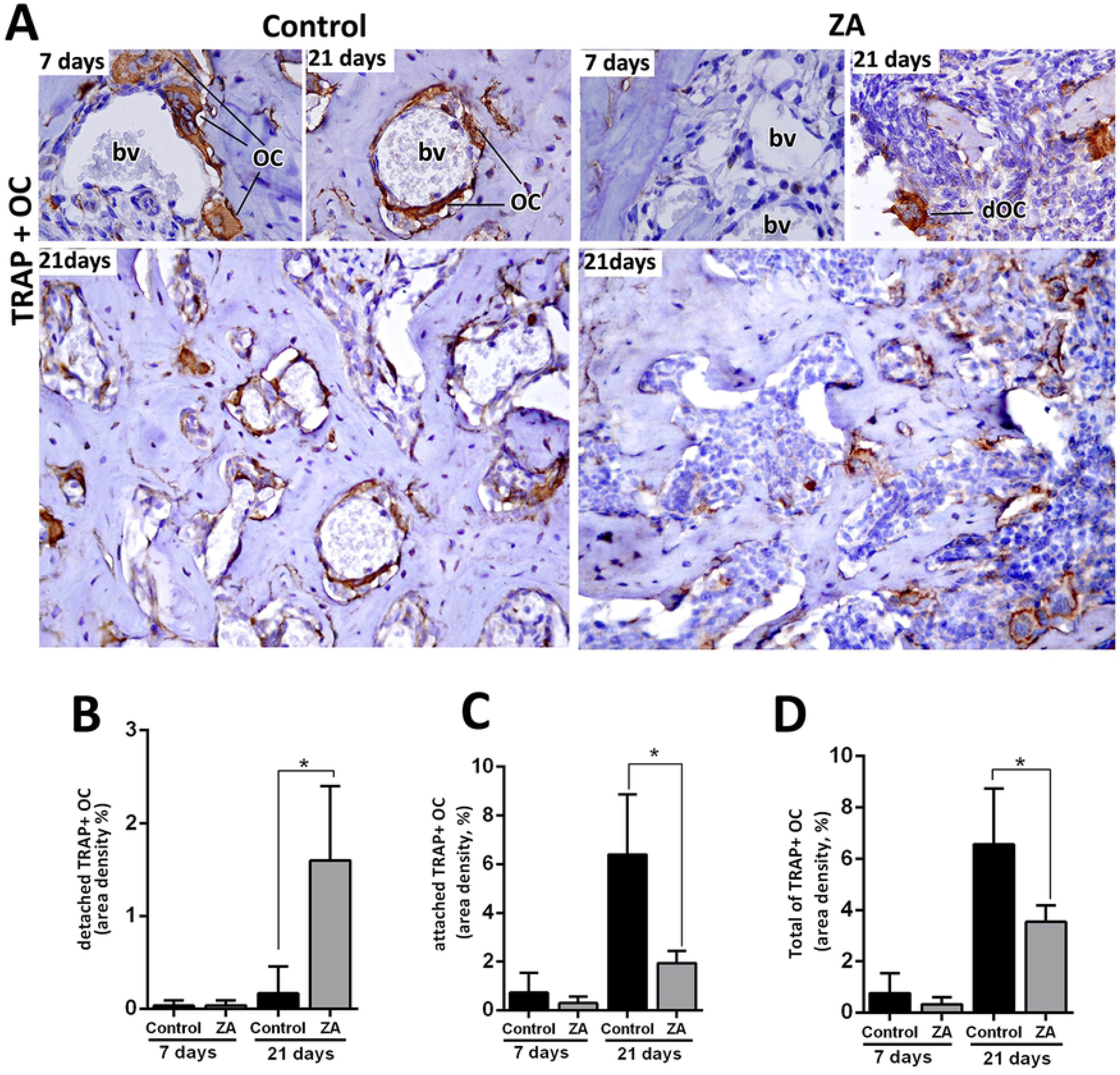
Immunolabeling and quantification of TRAP+OC in the Control and ZA treated mice. Senescent 129 Sv-WT femalemice received IP injections of 0.9% saline solution (Vehicle) or 250 µg/Kg one a week and upper right incisor were removed after 4 weeks of Vehicle or ZA treatments. Mice were euthanized for maxillary bones removal after 7days and 21days post tooth extraction.A) Representative transversal sections are observed throughout days 7 and 21, from the central area of alveolar sockets, for TRAP+OC in Control and ZA group. B-E) Quantitative and comparative analysis of detached TRAP+ OC (B), attached TRAP+ OC (C) and the Total of TRAP+OC considering the sum of detached and attached TRAP+ OC (D) in Control vs ZA treated mice at days 7 and 21 post-extraction. Results are presented as the means (±SD) of area density (%). Symbol *indicate a statistically significant difference vs control (p<0.05). DAB chromogen and counterstaining with Harris’ Hematoxylin.

**Table 1.** Summary of quantitative results for histomorphometry of area density (%)

| Parameters/Groups | Control 7d | ZA 7d | Control 21d | ZA 21d |
| --- | --- | --- | --- | --- |
| Blood clot | $3.27 \pm 3.10$ | $27.29 \pm 6.44^*$ | $0.00 \pm 0.00$ | $15.79 \pm 14.96^*$ |
| Inflam. Infiltrate | $5.53 \pm 1.17$ | $14.36 \pm 3.93^*$ | $1.62 \pm 0.62$ | $18.33 \pm 6.03^*$ |
| Osteocytes | $6.57 \pm 2.06$ | $1.63 \pm 1.32^*$ | $4.84 \pm 4.05$ | $2.13 \pm 2.88^*$ |
| Empty lacunae | $0.02 \pm 0.05$ | $0.75 \pm 0.95$ | $0.04 \pm 0.08$ | $7.43 \pm 7.85^*$ |
| Detached TRAP+OC | $0.03 \pm 0.05$ | $0.04 \pm 0.05$ | $0.016 \pm 0.28$ | $1.60 \pm 0.81^*$ |
| Attached TRAP+ OC | $0.72 \pm 0.82$ | $0.29 \pm 0.26$ | $6.40 \pm 2.45$ | $1.93 \pm 0.50^*$ |
| Total of TRAP +OC | $0.75 \pm 0.78$ | $0.32 \pm 0.27$ | $6.56 \pm 2.17$ | $3.53 \pm 0.64^*$ |
| TGFb+ cells | $13.34 \pm 6.67$ | $3.02 \pm 1.56^*$ | $4.16 \pm 1.58$ | $5.79 \pm 0.40$ |
| Runx-2+ cells | $10.71 \pm 1.63$ | $1.32 \pm 1.41^*$ | $8.93 \pm .76$ | $2.75 \pm 1.36^*$ |
| F4/80+ cells | $6.01 \pm 3.16$ | $1.90 \pm 1.71^*$ | $1.62 \pm 1.68$ | $3.42 \pm 1.61$ |

Results are presented as the means ( $\pm\text{SD}$ ) for each parameter.
Symbol \* indicate a statistically significant difference vs control at the same time point ( $p < 0.05$ ).

### TGF-β, Runx2, and F4/80+

Other important factors involved in healing events were also evaluated by immunostaining (Fig. 7A-D and S2 Fig). An increased level of expression for TGF-β, Runx2 and F4/80+ cells was detected in the initial stages of healing (7days) in Control group (Fig. 7A-E). Comparatively with ZA, the area density (%) of TGF-β, Runx2 and F4/80+ cells were increased in the Control considering the central area of alveolar sockets (Fig. 7 A-D). Also, the immunolabeling for PCNA+ epithelial cells in the oral mucosa, showed an intense nuclear fluorescence in the Control group basal layers, while PCNA+ cells were scarcely found in the same regions of the ZA group (S2 Fig.). At 21 days, Runx+2 cells were still found in increased number in the Control group compared to ZA. Mean and SD for all parameters are summarized in Table 1.

**Fig. 7.**
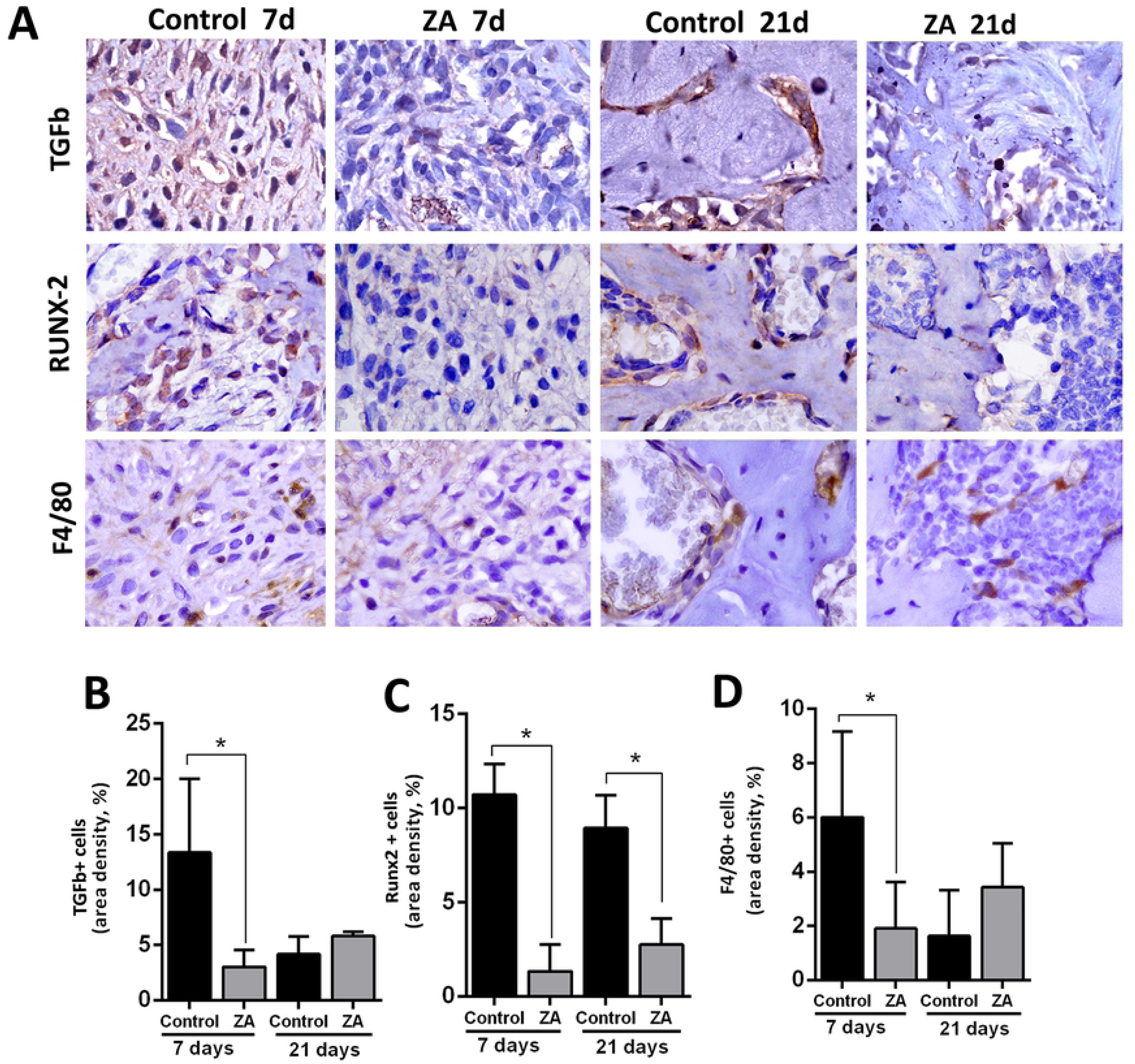
Immunolabeling and counting of TGFb, Runx-2 and F480+ cells of alveolar sockets post tooth extraction in Control and ZA treated mice. Senile 129 Sv-WT femalemice received IP injections of 0.9% saline solution (Vehicle) or 250 µg/Kg one a week and upper right incisor were removed after 4 weeks of Vehicle or ZA treatments. Mice were euthanized for maxillary bones removal after 7days and 21days post tooth extraction. A) Representative transversal sections are observed throughout days 7 and 21, from the central area of alveolar sockets, for TGFb, Runx2 and F4/80+ cells. B-D) Quantitative and comparative analysis TGFb+cells (B), Runx-2+ cells (C) and F4/80+ cells (D) in Control vs ZA treated mice at days 7 and 21 post-extraction. Results are presented as the means (±SD) of area density (%). Symbol *indicate a statistically significant difference vs control (p<0.05). DAB chromogen and counterstaining with Harris’ Hematoxylin.

## Discussion

This study investigated the bone healing after tooth extraction in senescent mice treated with zoledronic acid, as well as the usefulness of senescent 129/Sv female mice to understand the pathophysiology of MRONJ after tooth extraction. Previous MRONJ animal studies have not considered factors related to female human patients, such as age and post-menopause state, similar to what has been described in female human patients with risk factors for MRONJ [57]. Indeed, menopause female patients treated with nBPs are “10 fold higher risk to develop MRONJ at age 55 or older compared to young female patients” [1]. Matching the ages of laboratory mice and humans, the correlated age for mice used in this study (64-68 weeks) is about 50-52 years in humans [21] and the senescent changes in mice begin in middle age, around 40–60 weeks [58]. The average age of menopause in women is around 51 years [59] which match the age in 129Sv females who were found in a persistent diestrum and in the postmenopausal period the at 64-68 weeks old (Fig. 1A).

After demonstrating similar clinical preexisting conditions of MRONJ female patients in 129 Sv/female mice, MRONJ conditions were mimicked with upper right incisor extraction after 4 dosages of ZA or Vehicle (0.9% SS) (Fig. 1B), and following a previous alveolar bone repair model in mice [52, 53]. The Control group female mice presented a gradual mineralized and organic matrix deposition and maturation along 7 and 21 days (Fig. 2 and 3), compatible with other findings from 8 week-old C57Bl/6 WT mice alveolar sockets [52]. Also, histological features of bone healing and process of bone formation was gradual and consistent with other findings in control groups of mice [52, 53] and rats [60, 61], making the vehicle group suitable for comparison with ZA treated mice.

While for humans there are established concept and classification for MRONJ [1, 57, 62], animal MRONJ models are not comparable in terms of diagnosis and can vary according to the protocol applied to trigger MRONJ (S1 Table). In humans, MRONJ is defined as an area of exposed bone in the oral cavity that does not heal within 8 weeks, in a patient who has been receiving or has been exposed to antiresorptive (nBPs or denosumab) or antiangiogenic therapy, and has not had radiation therapy in the craniofacial region [2, 57]. Although bone exposure is a part of diagnosis criteria, the current staging system proposed by the AAOMS [1, 62] includes a variant of MRONJ without bone exposure (stage 0), with no clinical evidence of necrotic bone and/or infection. In our study, only 40% of ZA group animals clinically presented delayed epithelial socket closure compared to the Control (Fig. 1D) at 7 days, and a complete mucosal closure at 21 days. Additionally, no clinical suppuration and no biofilm evidences were noted in our model, because we do not use any periodontal pathogen inoculation. Indeed, laboratory mice (bred in ultra-hygienic environments) and human significantly differs in microbiota and pathogen-driven selection along their lives, which could reflect in different pattern of secondary infections in human and mice [20]. To address the oral bacterial influence on MRONJ pathogenesis, mice are inoculated with human periodontal pathogens in the local of oral injuries (e.g. *Fusobacterium nucleatum*) [26, 28].

In comparison to the available MRONJ mouse models (S1 Table), this study used a less traumatic triggering factor (tooth extraction of upper incisor instead of molars). Damage from extensive manipulation of bone, especially in advanced age conditions, does not reflect clinical conditions and could be a confounding factor for the healing process. In comparison with the present study, similar methodology was used in previous studies in rats (upper incisor extraction), in young (3 months old) [63] and senescent animals (20 months old) [61], where animals not necessarily developed clinical findings of MRONJ lesions, but all of them revealed histological features of this pathological conditions [63]. Despite of the differences in animal clinical findings with other studies [61, 63], 100% of our ZA samples presented delayed mineralized and organic bone matrix deposition, and also decreased level of collagen maturation along 7 and 21 days (Fig. 2 and 3). These findings are consistent with those in other studies using senescent female rats treated with ZA [60]. Col1a1 is significantly downregulated in MRONJ lesions in mice, affecting the collagen deposition, mainly by inhibition of TGF-β signaling pathway [23]. In our histopathological analysis, 100% of our ZA treated mice presented a delayed bone healing and MRONJ lesions histological characteristics (Fig. 4 and 5). These compare with other findings in mice [23, 39], rats [60, 61, 63] and humans [12, 13], including intense leukocyte infiltration, areas of non-vital bone and discrete bone sequestrum.

To accurately analyze the effect of ZA administration on bone cells, we investigated the morphological and histomorphometric changes in osteocytes and empty lacunae (Fig. 3D-E) as well as TRAP+ OC’s (Fig. 6). As expected, ZA group presented a decreased number of Total TRAP+ OC’s compared to the Control, as observed in other animal models [60] and humans [64]. The majority of TRAP+ OC’s in ZA samples were detached from the bone matrix which resembled MRONJ characteristics in senescent rats [61] and human biopsies [64]. Additionally, an increased quantity of empty lacunae was found in the ZA E site compared to Control E site (Fig. 4). Empty lacunae is one diagnostic factor for osteonecrosis [12, 13] and is consistent with other findings in mice MRONJ lesions [23]. Interestingly, the greatest amount of empty lacunae was found at E sites, but not in the NE sites in ZA-treated mice (S1 Fig.). This finding highlights a hindrance to bone repair processes in which, under normal circumstances, osteocytes undergo apoptosis and recruit osteoclasts to resorb damaged bone [67]. Furthermore, it emphasizes that trauma is crucial for osteocyte apoptosis, which is a hallmark characteristic of necrotic bone [12, 13].

Both osteoclast and osteocyte biology may assist in understanding the pathogenesis of MRONJ, partially because of the osteocyte signaling to osteoclasts in injury and under mechanical loading conditions [65]. At the ultrastructural level, nBP’s target OC’s by blocking the mevalonate pathway, thereby negatively interferes with the organization of their cytoskeleton, thus blocking the formation of the ruffled border [66]. The cell resorption compartment, where the protons and proteolitic enzymes are released for the dissolution of bone mineral and organic contends, cannot be formed leading to apoptosis of OCs [47].

While nBP’s induce OC apoptosis, some evidence has revealed an opposite effect on the osteocytes, which is a required effect for blocking bone loss in osteoporotic long bones [67, 68]. However, osteocyte apoptosis is essential to replace damaged bone [67]. When osteocytes age, some of these cells die, though in areas of low bone turnover, such as ear bones, empty lacunae do not necessarily indicate necrotic bone [71]. In aged bone, senescent osteocytes will actually increase their number of dendritic processes [69, 77], indicating increases in regulation of bone homeostasis.

One important function of osteocytes is to sense mechanical pressure on bone and send signals to resorb bone where deemed appropriate [73–74]. A previous study by Marx in 2014 reported that the incidence of MRONJ in the oral cavity is highest in areas of high mechanical load on bone (posterior mandibular lingual cortex, edentulous alveolar ridge covered by dentures, and the lamina dura). Marx also reported that mechanical loading of bone causes decreased osteocyte OPG secretion, thus an increased RANKL/OPG ratio and increased osteoclast recruitment to resorb the involved bone [72]. Furthermore, excessive or insufficient levels of mechanical loading cause apoptosis of osteocytes and induction of osteoclastic bone resorption [73–75]. If this bone is nBP-laden, these resorbing osteoclasts would undergo apoptosis and leave the remaining bone with its empty lacunae. When tooth extraction causes bony injury to the alveolar socket, the healing process is similar to aforementioned bone resorption. Most osteocytes more than 1-2 mm from an bone fracture do not die [74] and will increase RANKL and decrease OPG production [76]. Concurrently, osteocytes within 1-2 mm of the fracture site undergo apoptosis and release RANKL-wielding vesicles that promote osteoclast recruitment [77]. Osteocyte apoptosis in damaged bone therefore leaves an acellular bone matrix ready to be resorbed by recruited osteoclasts [74]. The MRONJ phenotype results in part from the detachment and death of recruited osteoclasts when they ingest nBP-laden bone. Furthermore, the bone matrix has empty osteocyte lacunae from the initial response to the trauma, as observed in the injured socket areas in this study (S Fig. 1 and Fig. 3).

Our data has demonstrated a decreased number of TGF-β+ cells and Runx-2+ cells (committed osteoblasts) in ZA treated mice, as similarly observed in other MRONJ mice [23] and rat models [61], respectively. TGF-ß1 signaling pathway is necessary to activate the transcription factor Runx-2 (a global regulator of osteogenesis) (Lee 2000) and both are significantly up-regulated in control conditions along alveolar bone repair in mice, concomitant with new bone differentiation and bone deposition [52, 53]. In this context, another important effect of nBP’s is on new bone cells’ differentiation in injured sites, since TGF-β1 signaling pathway is also significantly suppressed in MRONJ lesions at the clinical level [78].

Finally, although nBP influence on bone cells has been previously demonstrated, its action on immune cells remains largely unclear. Several studies have revealed that monocytes and macrophages are capable of internalizing nBP’s [66, 79]. Post-tooth extraction experimental models performed in mice revealed a high polarization of M1 macrophages induced by IL-17 increase in non-healing sockets. This evidence demonstrates an imbalance in M1/M2 ratio, which is related to the pathogenesis of disease [17]. In this study, a decreased migration of F4/80+ cells at 7 days was observed in ZA mice compared to the Control, while a slight increase in migration was observed at 21 days in MRONJ lesions (Fig.7-E). Accordingly, a delayed removal of blood clot and an increased persistence of general inflammatory infiltration were also noted in ZA group compared to the Control (Fig. 3BC). F4/80+ cells are supposed to be mainly macrophages, as has been demonstrated in similar tooth extraction mouse model [53]. In this context, macrophages are supposed to be important for producing growth factors, activation of immune response and clearance of tissue debris, blood clot and necrotic cells [53]. According to these findings, this study also draws the attention for future evaluations of interferences on immunological system caused by the nBPs.

## Conclusions

In summary, despite the limitations of this study, the presented findings are consistent with the accepted MRONJ phenotype, that include delayed blood clot and debris removal, increased and disruption of inflammatory infiltration, increased occurrence of empty lacunae and bone sequestrum in injured alveolar socket, fewer TRAP+ OCs, and potential inhibition of bone-related growth factors accompanied by delayed bone matrix formation and maturation (Fig. 8). Considering all these observations and comparing with previous descriptions of MRONJ in humans, this MRONJ mouse model offers a novel experimental tool for development of future pre-clinical interventions.

**Fig. 8.**
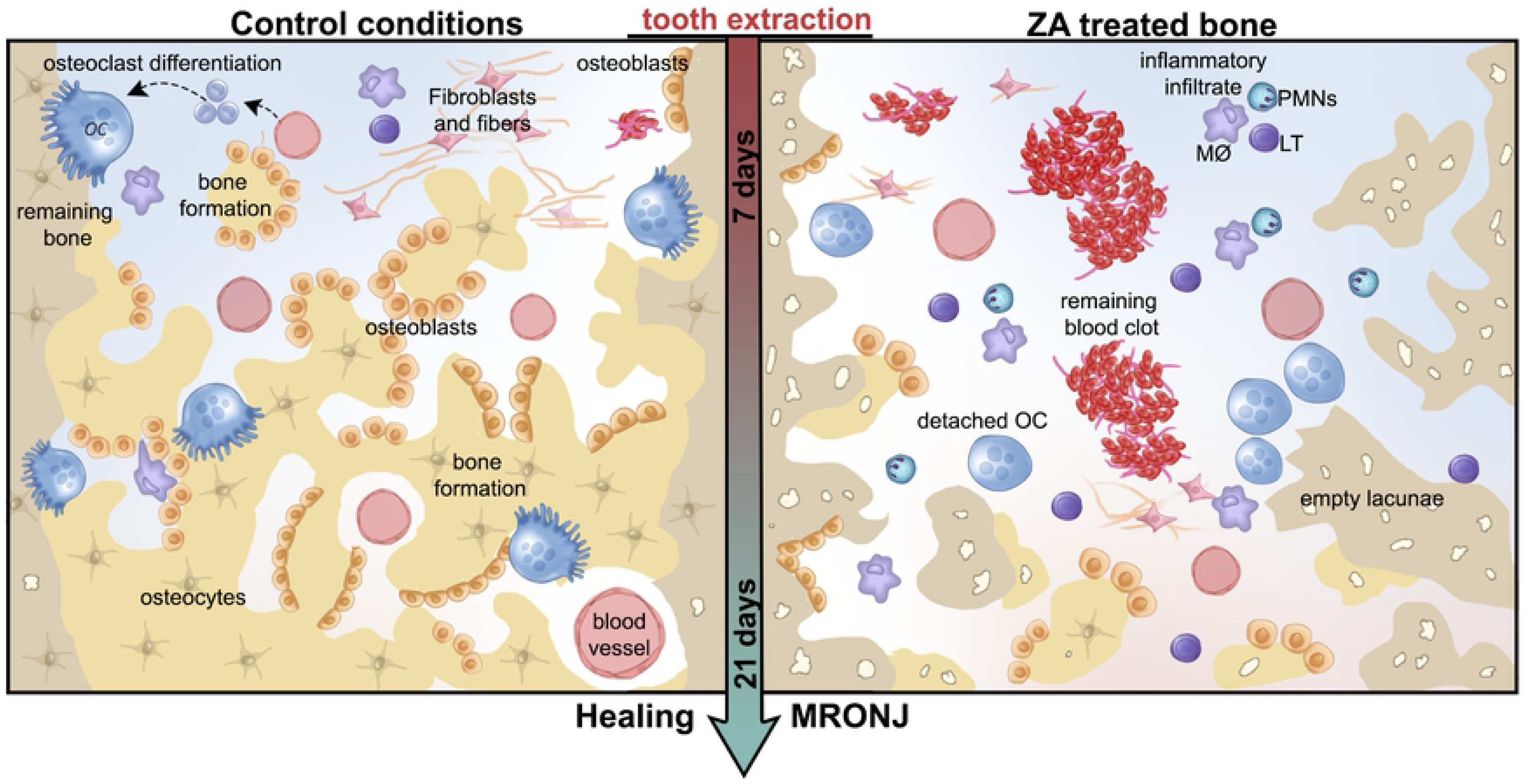
Graphical representation of microscopic events during initial and late time points post tooth extraction in Control conditions compared to MRONJ model in senescent 129 Sv WT female mice. In summary, senescent 129 Sv female mice treated with cumulative dosages of Zoledronic acid present a disrupted healing process post tooth extraction compared to Control conditions. While bone formation and remodeling occurs along 21 days in control conditions, in the MRONJ phenotype, a delayed blood clot and debris removal, increased inflammatory infiltration, increased occurrence of empty lacunae and bone sequestrum, detached osteoclasts are seen until the later time points of 21 days.

## Acknowledgements

The authors acknowledge Ms. Maira Cristina Couto for her excellent technical assistance in this study.

## Supporting Information

**S1 Table. Summary of MRONJ mice models**

**S1 Fig.** Representative transversal sections are observed throughout 7 days from alveolar bone of extraction sites (1-4) and control sites (5-8) in Control group (A) and ZA (B) group. A) Control group extraction sites (A1-A4) and Control sites (A5-A8) filled with osteocytes (black arrows). B) ZA group extraction sites (A1-A4) present several empty lacunaes (red arrows), while Control sites (A5-A8) remain filled with osteocytes (black arrows). Histological slides were stained with H&E (upper panel) and GT+Alcian blue (lower panel) and images were captured at 2x (entire section at left side) and 100x magnification (panels).

**S2 Fig. Immunolabeling for PCNA+ epithelial cells from the alveolar socket area at 7days post tooth extraction in senile 129 Sv-WT female.** Mice received IP injections of 0.9% saline solution (Vehicle) or 250 µg/Kg one a week and upper right incisor were removed after 4 weeks of Vehicle or ZA treatments. Mice were euthanized for maxillary bones removal after 7days post tooth extraction. Secondary antibody Cy3 (#715-165-150, Jackson ImmunoResearch Laboratories, West Grove, PA, USA) for detection of PCNA and DAPI (D9542-50, Sigma-Aldrich Corp., St. Louis, MO, USA) for nuclear staining.

